# Multifunctional flexible electro-optical arrays for simultaneous spatiotemporal cardiac mapping and modulation

**DOI:** 10.1101/2022.05.14.491979

**Authors:** Sofian N. Obaid, Zhiyuan Chen, Micah Madrid, Zexu Lin, Jinbi Tian, Camille Humphreys, Jillian Adams, Nicolas Daza, Jade Balansag, Igor R. Efimov, Luyao Lu

## Abstract

Bioelectronic devices that allow simultaneous accurate monitoring and control of the spatiotemporal patterns of cardiac activity provide an effective means to understand the mechanisms and optimize therapeutic strategies for heart disease. Optogenetics is a promising technology for cardiac research due to its advantages such as cell-type selectivity and high space-time resolution, but its efficacy is limited by the insufficient number of modulation channels and lack of simultaneous spatiotemporal mapping capabilities in current cardiac optogenetics tools. Here we present soft implantable electro-optical cardiac devices integrating multilayered highly uniform arrays of transparent microelectrodes and multicolor micro-light-emitting-diodes in thin, flexible platforms for mechanically compliant high-content electrical mapping and single-/multi-site optogenetics and electrical stimulation without light-induced artifacts. Systematic benchtop characterizations, together with *ex vivo* and *in vivo* evaluations on healthy and diseased small animal and human hearts demonstrate their functionalities in real-time spatiotemporal mapping and control of cardiac rhythm and function, with broad applications in basic and ultimately clinical cardiology.

## 1. Introduction

Heart disease has a significant negative impact on the quality of life and is the leading cause of mortality in the world.^[1–2]^ Heart disease arises from asynchrony and abnormalities in the complex and coordinated electro-mechanical properties over time and space. Therefore, devices that allow monitoring and controlling of the spatiotemporal dynamics of cardiac activity are crucial for unraveling the pathophysiology of heart disease and developing effective treatment therapies in clinical cardiology practice. Implantable electronic pacemakers play an essential role in treating various types of arrhythmias and heart failure and studying cardiac physiology by changing the membrane potential and triggering an action potential with an electric current.^[3–4]^ However, electrical stimulation can lead to adverse effects on cell health and integrity due to the cell membrane electroporation and redox processes.^[5–6]^ The electrical fields created by the stimulation electrodes will generate electrical crosstalk between stimulation and recording electrodes and result in recording artifacts.^[7]^ Furthermore, electrical stimulation is unable to target specific subtypes of cardiac cells. Optogenetics uses light to modulate the activity of genetically targeted cell types through photosensitive ion channels and pumps.^[8]^ Despite its initial use in neuroscience research to control neural circuits,^[8–9]^ optogenetics has now been applied as a promising tool in cardiology for pain-free, low-energy optical pacing and defibrillation with cell-type specificity.^[10–12]^ In addition, cardiac optogenetics generally interferes less with simultaneous electrical readout of cardiac activity compared to electrical stimulation. Currently, cardiac optogenetics is primarily used in *in vitro* and *ex vivo* cardiac studies with cell cultures or explanted perfused hearts.^[13–16]^ More *in vivo* research is crucially needed to fully exploit the unique opportunities cardiac optogenetics offers for mechanistic investigations of heart function in health and disease.

Small animal models such as mice and rats are the main rodent species that could be genetically engineered for *in vivo* cardiac optogenetics research and their heart electrophysiology is a good approximation of the human electrophysiology.^[17]^ Implantable cardiac devices that combine precise light delivery to targeted heart regions of small animals with electrophysiological readout capabilities remain a major technological challenge. State-of-the-art tools primarily rely on optical fibers connected to external light sources for single site monochromatic optogenetics modulation with physically separated body surface electrocardiographic (ECG) electrodes for recording the resulting cardiac electrical activity.^[18–20]^ However, those devices lack the spatial control of optogenetics modulation patterns, which prevent the fine perturbation of the electrophysiological state and the evaluation of multisite cardiac pacing strategies in terminating arrhythmias and heart failure.^[21]^ The monochromatic feature limits the capability to simultaneously manipulate more than one cell type. In addition, although surface ECG measurement is the current gold standard for monitoring heart rhythm, it does not allow direct probing of the spatiotemporal patterns of regional and tissue level cardiac activity, such as cardiac excitation waves and conduction system on the epicardial surface, which are central to heart function and different in health and disease.^[22–23]^ In this context, a highly integrated, soft, implantable cardiac device that seamlessly interfaces with multiple locations on the curvilinear epicardium for spatiotemporal mapping and optogenetics control of the dynamic cardiac electrophysiological behavior would be indispensable, particularly for studying and controlling the complex, lift-threatening, space-time cardiac events like arrhythmias and heart failure.

Recently we demonstrated an innovative bioelectronic device design by integrating a transparent metal grid microelectrode ontop of a microscale light source for precise, colocalized single site cardiac optogenetics and electrical recording.^[24]^ In this work, we adopt this approach and take a major leap forward by creating soft, flexible, multifunctional, highly uniform hybrid electro-optical array technologies that contain a 16-channel optically transparent gold (Au) grid microelectrode array (MEA) and a 4-channel multicolor micro-light-emitting-diode (μ-LED) array in multilayer stacked architectures. The microelectrodes exhibit a high optical transparency across a broad spectral region from 400 nm to 800 nm and allow light from the underneath multicolor μ-LEDs to pass through the microelectrodes for colocalized site-specific optogenetics modulation and electrical mapping with negligible light-induced artifacts. Systematic electrical, electrochemical, optical, mechanical, and thermal characterizations illustrate the robustness and uniformity of the MEA and μ-LED array. *Ex vivo* and *in vivo* validations by crosstalk-free spatiotemporal electrical mapping and optogenetics pacing of mouse and human hearts under different conditions, such as sinus rhythm and ventricular tachycardia (VT) illustrate the function, form factor, reliability, and capability of the electro-optical arrays in both monitoring and cell-specific managing cardiac rhythm (e.g., direction and velocity of excitation waves), in ways that are impossible with conventional techniques.

## 2. Results and Discussion

### 2.1. Design and Fabrication of the Electro-optical Array Devices

**Figure 1**a presents an exploded schematic illustration of the electro-optical array device containing a 4 × 4 transparent Au grid MEA and a 2 × 2 μ-LED array. Each μ-LED site has a transparent microelectrode directly on top. The centers of the colocalized microelectrode and μ-LED are well aligned. *Ex vivo* optical mapping results reveal that the emission power from a blue μ-LED in the electro-optical array device decreases exponentially to <1% of the initial value after travelling 3 mm on the epicardial surface of a mouse heart (Figure 1b). As a result, we use 3 mm as the pitch value for the μ-LED array to avoid optical cross-excitation, where light from one μ-LED excites or inhibits the activity of cardiomyocytes at a nearby site. Meanwhile, the pitch value for the MEA is 1 mm to achieve a high spatial resolution electrical mapping of the cardiac signal propagation within the region fully covered by the μ-LED array. The numbers of μ-LEDs and microelectrodes in the devices are sufficient to perform single-chamber, dual-chamber (atrial and ventricular) or biventricular pacing while the overall device dimension (3 × 3 mm^2^) is on par with the sizes of ventricles of small animal models.^[25–26]^ Device fabrication begins with spin-coating of a SU-8 planarization layer (thickness, 7 μm) onto a flexible, transparent, and biocompatible polyethylene terephthalate (PET) substrate (thickness, 25 μm). Lithographically patterned bilayer chromium/copper (Cr/Cu, thickness, 20/100 nm) lines serve as the electrical interconnects for the μ-LEDs. Micro-transfer printing with polydimethylsiloxane (PDMS) microstamps places the 2 × 2 μ-LED array onto the desired locations on the Cr/Cu interconnects. A reflow soldering process electrically and mechanically joins the μ-LEDs with the interconnects. A SU-8 layer (thickness, 60 μm) insulates the μ-LED array. Afterward, the 4 × 4 Au grid (thickness, 40 nm) transparent MEA and interconnects are lithographically patterned on the SU-8 passivation layer. Another SU-8 layer (thickness, 7 μm) encapsulates the device, defines the MEA sizes, and completes the fabrication. Detailed information of device fabrication appears in Figure S1 (Supporting Information) and the Experimental Section.

**Figure 1.**
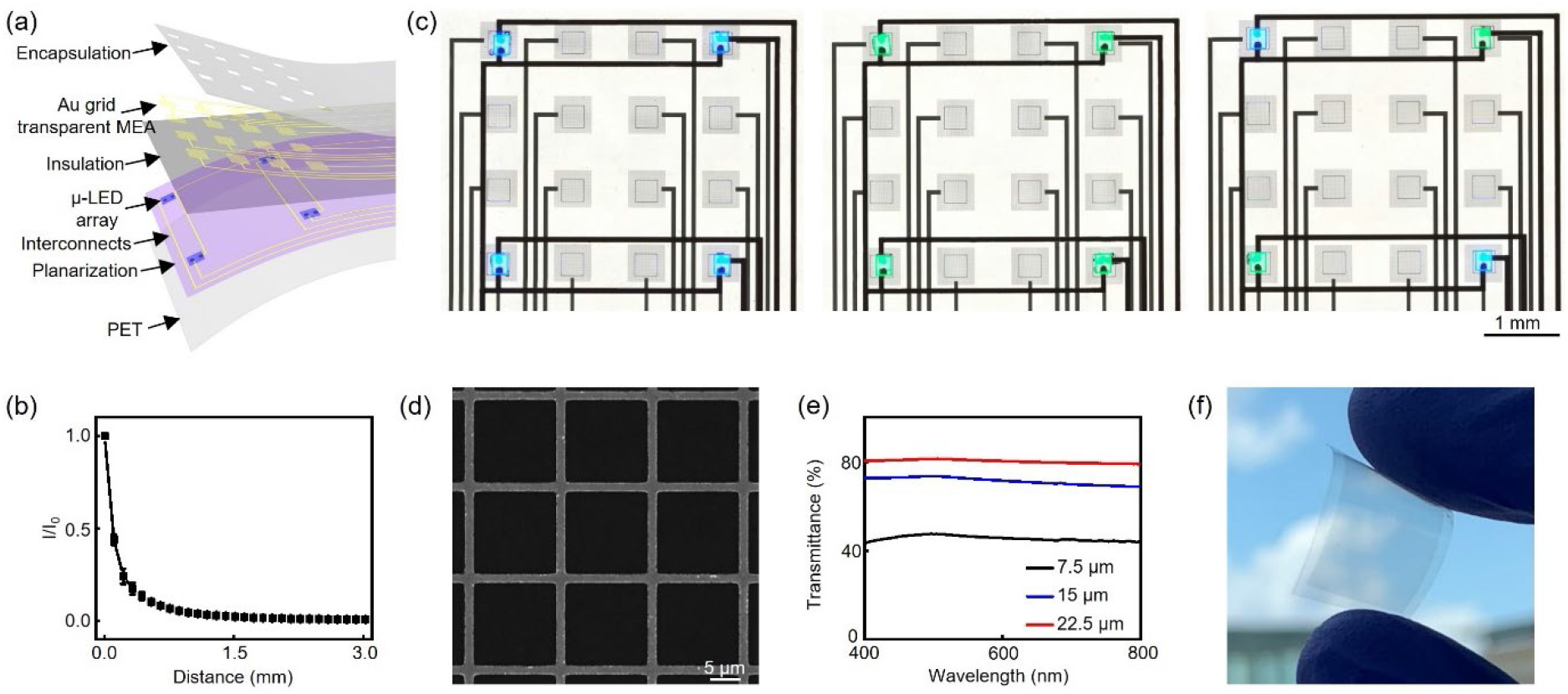
Multifunctional electro-optical array devices for simultaneous electrical mapping/pacing and optogenetics modulation of cardiac activity. a) Exploded illustration of the electro-optical array device consisting of a 4 × 4 Au grid transparent MEA and a 2 × 2 μ-LED array. b) Attenuation of optical irradiance as a function of distance away from a blue μ-LED in the electro-optical array device on the surface of a mouse heart. I is the irradiance on the epicardial surface at a certain distance away from the blue μ-LED site, I_0_ is the initial irradiance at the blue μ-LED site. c) Optical images of the fully integrated multilayered electro-optical array devices. From left to right: devices with 4 blue μ-LEDs on, 4 green μ-LEDs on, 2 blue and 2 green μ-LEDs on. d) Scanning electron microscopy image of the Au grid network in (c). The line width of the individual grid is 1.5 μm and the pitch value between different lines in the grid network is 15 μm, respectively. e) Transmittance spectra of Au grids with a 1.5 μm grid width and different pitch values at 7.5 μm, 15 μm, and 22.5 μm, respectively. f) Optical image of a 1 × 2 cm^2^ transparent Au grid network on a flexible PET film.

Channelrhodopsin-2 (ChR2) is one of the most widely used blue light activated opsins in optogenetics.^[10]^ Meanwhile, opsins with red-shifted excitation spectra are of interest due to their increased penetration depth in the tissue and associated advantages to capture a large volume of tissue deep within the myocardium. Our fabrication strategy is versatile and allows the integration of multicolor μ-LEDs to satisfy different needs in the community. As a proof-of-concept demonstration, Figure 1c presents electro-optical array devices with 4 blue, 4 green, interdigitated blue and green μ-LEDs. The blue and green μ-LEDs exhibit emission maxima at 462 nm and 531 nm, respectively (Figure S2, Supporting Information). Those devices enable parametric optogenetics control of two cardiac cell types (e.g., when excitatory and inhibitory opsins are co-expressed) and mapping of the resulting cardiac activity. The μ-LEDs are connected in parallel for precise control of single site, multisite, and multicolor cardiac optogenetics modulation parameters (Figures S3, S4, Supporting Information). Each microelectrode consists of a Au grid network with the grid width and pitch values of 1.5 μm and 15 μm (Figure 1d), respectively. The grid parameters can be tuned to adjust the physical properties of the microelectrodes. Figure 1e shows that the average transmittance values of the Au grid network (over 5 devices) at 550 nm increase from 46.4 ± 0.43% to 72.5 ± 0.93% and 80.7 ± 0.97% with the grid pitch values increase from 7.5 μm to 15 μm and 22.5 μm, respectively. Importantly, our lithography-based fabrication method allows scaling up the overall dimensions of the MEA. Figure 1f displays a 1 × 2 cm^2^ transparent Au grid/PET film. Meanwhile, the μ-LEDs can be designed into larger-area and higher-density arrays using automated mass micro-transfer printing techniques.^[27–29]^

### 2.2. Performance of the Electro-optical Array Devices

Electrochemical impedance spectroscopy (EIS) and cyclic voltammetry (CV) measurements evaluate the electrical recording and stimulation capabilities of the Au grid transparent MEAs. **Figure 2**a presents the EIS results of the Au grid microelectrodes with varying pitch values measured at a frequency range from 100 Hz to 10 kHz in a 1× phosphate-buffered saline (PBS) solution. The microelectrodes have dimensions of 300 × 300 μm^2^ unless otherwise noted. The impedance values at 1 kHz increase from 6.50 kΩ to 12.0 kΩ, and further to 18.5 kΩ with pitch values increasing from 7.5 μm, to 15 μm and 22.5 μm, respectively. The normalized impedance of the Au grid microelectrodes is comparable to previously reported flexible transparent microelectrodes at similar transmittance values, such as graphene, carbon nanotube, metal nanomesh, etc.^[30–36]^ The dimensions of each microelectrodes in the electro-optical array device can be reduced to 50 × 50 μm^2^ (Figure S5, Supporting Information), comparable to individual cardiomyocytes (~20 × 100 μm^2^),^[37–38]^ while still exhibit a moderate impedance value of 269 ± 10.2 kΩ (Figure S6, Supporting Information). This will allow tissue level cardiac electrical mapping using the MEA with cellular resolution.

**Figure 2.**
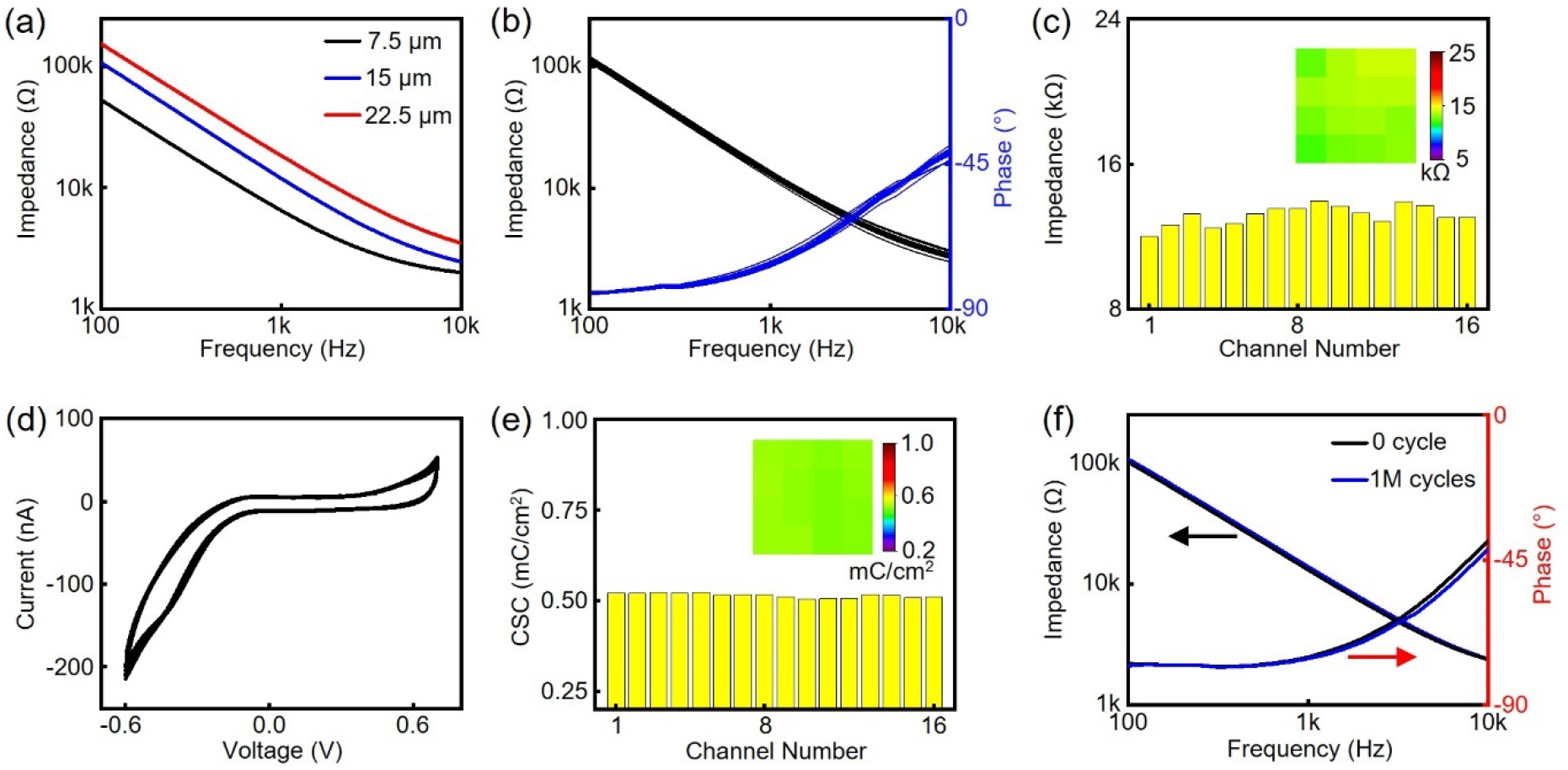
Electrochemical characterizations of the Au grid transparent MEAs. a) Impedance spectra of Au grid microelectrodes with a 1.5 μm grid width and different pitch values at 7.5 μm, 15 μm, and 22.5 μm, respectively. b) Impedance and phase spectra of the 16 microelectrodes in a Au grid transparent MEA. c) Impedance histogram of the 16-channel Au grid transparent MEA in (b). Inset: Impedance colormap of the microelectrodes with respect to actual microelectrode position in the MEA. d) CV spectra of the 16 microelectrodes in a Au grid transparent MEA. Scan rate: 100 mV/s. e) CSC histogram of the 16-channel Au grid transparent MEA in (d). Inset: CSC colormap of the microelectrodes with respect to actual microelectrode position in the MEA. f) Impedance and phase spectra of the Au grid microelectrodes before and after 1 million cycles of electrical stimulation at 8 Hz, 1.6% duty cycle, and 0.5 V.

The MEA exhibits a high degree of uniformity with an average impedance and phase angle of 13.2 ± 0.56 kΩ and −76.1 ± 0.71° at 1 kHz (Figure 2b,c). The phase angle suggests a nearly capacitive interface of the Au grid microelectrodes. The average impedance and phase angle over 5 multifunctional devices (80 channels in total) are 13.8 ± 2.54 kΩ and −75.0 ± 1.74°, respectively. The highly uniform, reproducible, and superior electrochemical performance is crucial for low noise mapping of the cardiac electrophysiological signals. Charge storage capacity (CSC) is an important parameter to determine the electrical stimulation capability of a microelectrode. Figure 2d,e presents the CV results from −0.60 V to 0.70 V and extracted CSC values of all 16 MEA channels. This voltage range is chosen to fall within the water hydrolysis window. The average CSC value is 0.51 ± 0.01 mC/cm^2^, respectively. The impedance and phase spectra of the Au grid microelectrodes remain stable after 1 million cycles of continuous electrical stimulation at 8 Hz and 0.5 V in a PBS solution (Figure 2f and Figure S7, Supporting Information). Those results demonstrate the robustness of the microelectrodes for continuous, repeated, chronic cardiac electrical interfacing.

The high-fidelity electrical recording capability is further demonstrated by recording a sine wave input (10 Hz, 20 mV peak-to-peak amplitude) delivered to a PBS solution by a platinum electrode. **Figure 3**a presents the recorded sine wave with no observable decrease in signal amplitude from a Au grid microelectrode. The power spectral density (PSD) results (Figure 3b) provide detailed frequency domain information on the recorded signals. The large peak corresponds to the 10 Hz input signal. Figure 3c and Figure S8 (Supporting Information) show the highly uniform signal-to-noise ratio (SNR) and root-mean-square (RMS) noise across all 16 microelectrode channels. The average SNR and RMS noise are 40.3 ± 0.28 dB and 41.7 ± 4.09 μV, respectively. More recording results with sine wave inputs at different cardiac related frequencies (from 5 Hz to 8 Hz) are summarized in Figure S9 (Supporting Information). A superior mechanical flexibility is important to form a conformal contact between the implantable devices and the curvilinear heart surface. The MEA (Figure 3d) and μ-LED array (Figure 3e) maintain stable electrochemical and optoelectronic performance after 5,000 bending cycles against a small radius of 5 mm, comparable to the anatomical features of interest in mouse hearts. Soaking test in a PBS solution (pH 7.4) at 37 °C reveals negligible impedance degradation from the microelectrodes (Figure 3f) and minimal changes in the current-voltage characteristics of the μ-LEDs (Figure 3g) for up to 4 weeks. Those results suggest the suitability of the electro-optical array devices for long-term applications. Figure 3h summarizes the temperature increases on the surface of the blue μ-LED in air as a function of duty cycle with a frequency of 7 Hz and an output irradiance of 25.3 mW/mm^2^, relevant to the μ-LED operating parameters used in optogenetics. The maximum temperature increase is within 0.4 °C for duty cycles ≤5%, which falls well below the thermal thresholds for cardiac tissue damage.^[39]^

**Figure 3.**
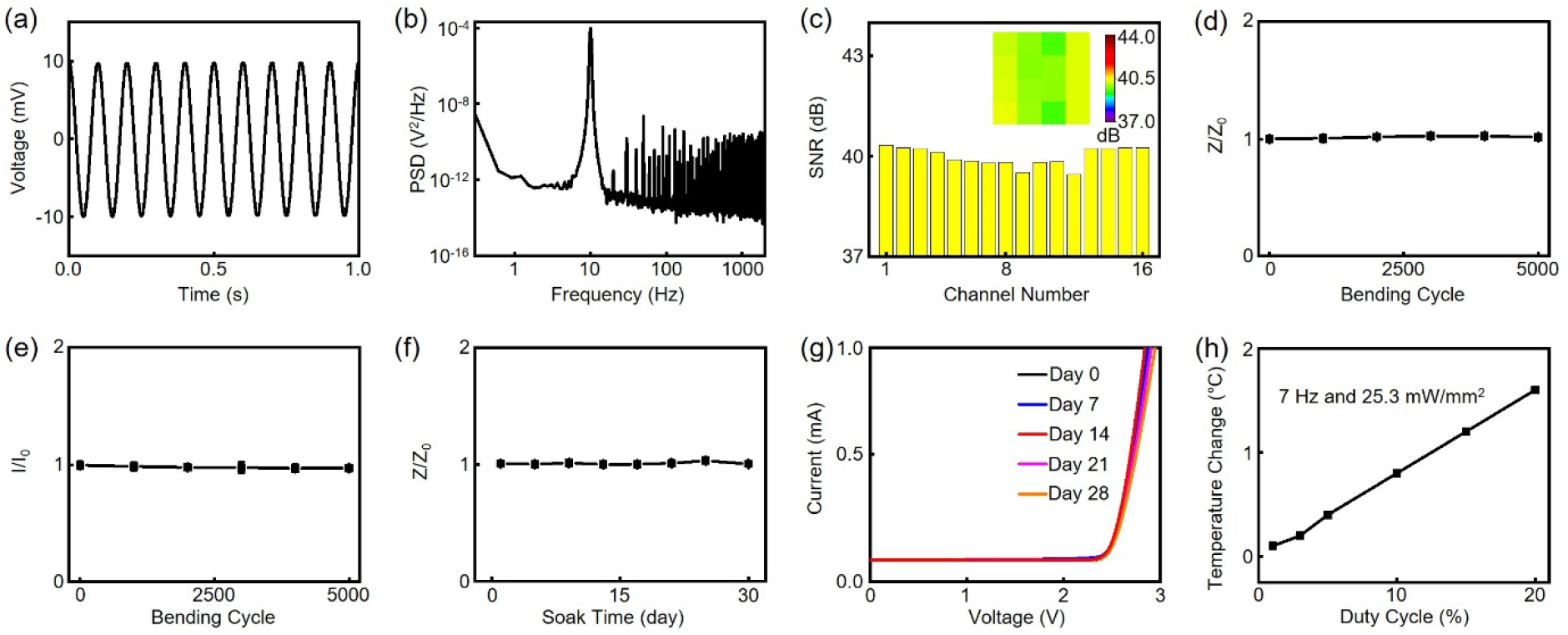
Benchtop measurements of the electro-optical array devices. a) Representative benchtop electrical recording output of a 10 Hz, 20 mV peak-to-peak input sine wave in a PBS solution by the Au grid microelectrode. b) Representative PSD graph of the recorded sine wave output in (a). c) SNR histogram of the 16 microelectrodes in a Au grid transparent MEA from the benchtop electrical recording in (a). Inset: SNR colormap of the microelectrodes with respect to actual microelectrode position in the MEA. Au grid microelectrode impedance (d) and μ-LED irradiance (e) as a function of bending cycle with a bending radius of 5 mm. Z and I are the impedance and irradiance at a specific bending cycle, Z_0_ and I_0_ are the initial impedance and irradiance, respectively. Changes of impedance from the Au grid microelectrodes (f) and current-voltage characteristics from the μ-LEDs (g) in the electro-optical array devices as a function of soaking time in a PBS solution at 37 °C. h) Temperature change versus duty cycle from the blue μ-LEDs in an electro-optical array device at 7 Hz and 25.3 mW/mm^2^ output irradiance.

### 2.3. *Ex Vivo* Demonstrations of Cardiac Electrical Mapping and Pacing

Experiments on *ex vivo* Langendorff-perfused mouse hearts first validate the electrical mapping and pacing functions of the electro-optical array devices. The device is attached onto the epicardial surface of a mouse left ventricle (LV). **Figure 4**a shows the representative electrical and optical activation maps during sinus rhythm and the sequence of the 16 microelectrode channels in the devices. The optical voltage fluorescence signals are obtained with a complementary metal-oxide-semiconductor (CMOS) imaging camera. The electrical and optical mapping signals show strong correlations, suggesting the high-fidelity mapping capabilitiy of the MEA. Here the optical signals represent average activity over ~0.5 mm in depth of the heart tissue and the electro-optical array device collects cardiac wave propagation from the epicardium.^[40]^ Moreover, we can achieve electrical pacing using one or more microelectrodes in the MEA and simultaneously evaluate the pacing effects by electrical mapping from the surrounding microelectrodes. This will enable potential comparison between electrical and optogenetics pacing effects. Figure 4b shows such a demonstration, where microelectrode channel 2 stimulates the LV at 5 Hz, 1% duty cycle, and 2 V while the other 15 microelectrode channels map the resulting propagation of cardiac electrical waves. As expected, cardiac activation originates at the electrical pacing site and the electrical activation map matches well with the optical one. The wave propagation can be precisely manipulated by the location (Figure S10, Supporting Information) and number (Figure S11, Supporting Information) of microelectrodes used in electrical pacing. Electrical strength duration curves (n=3) from microelectrodes at the corner and inner locations in the MEA (Figure 4c and Figure S12, Supporting Information) show that the required thresholds and pulse widths to achieve successful pacing are comparable between the Au grid transparent MEA and an external platinum-iridium (Pt-Ir) reference electrode. Moreover, we show that the devices can record electrogram (EG) signals from a human ventricular tissue slice (Figure 4d and Figure S13, Supporting Information). Here, the MEA measured transverse conduction velocity from the tissue slice is 24.6 cm/s, similar to the value obtained from optical mapping (24.8 cm/s). These results highlight the capabilities of the electro-optical array devices in multifunctional electrical mapping and pacing of different cardiac tissue models and the compatibility with optical mapping techniques.

**Figure 4.**
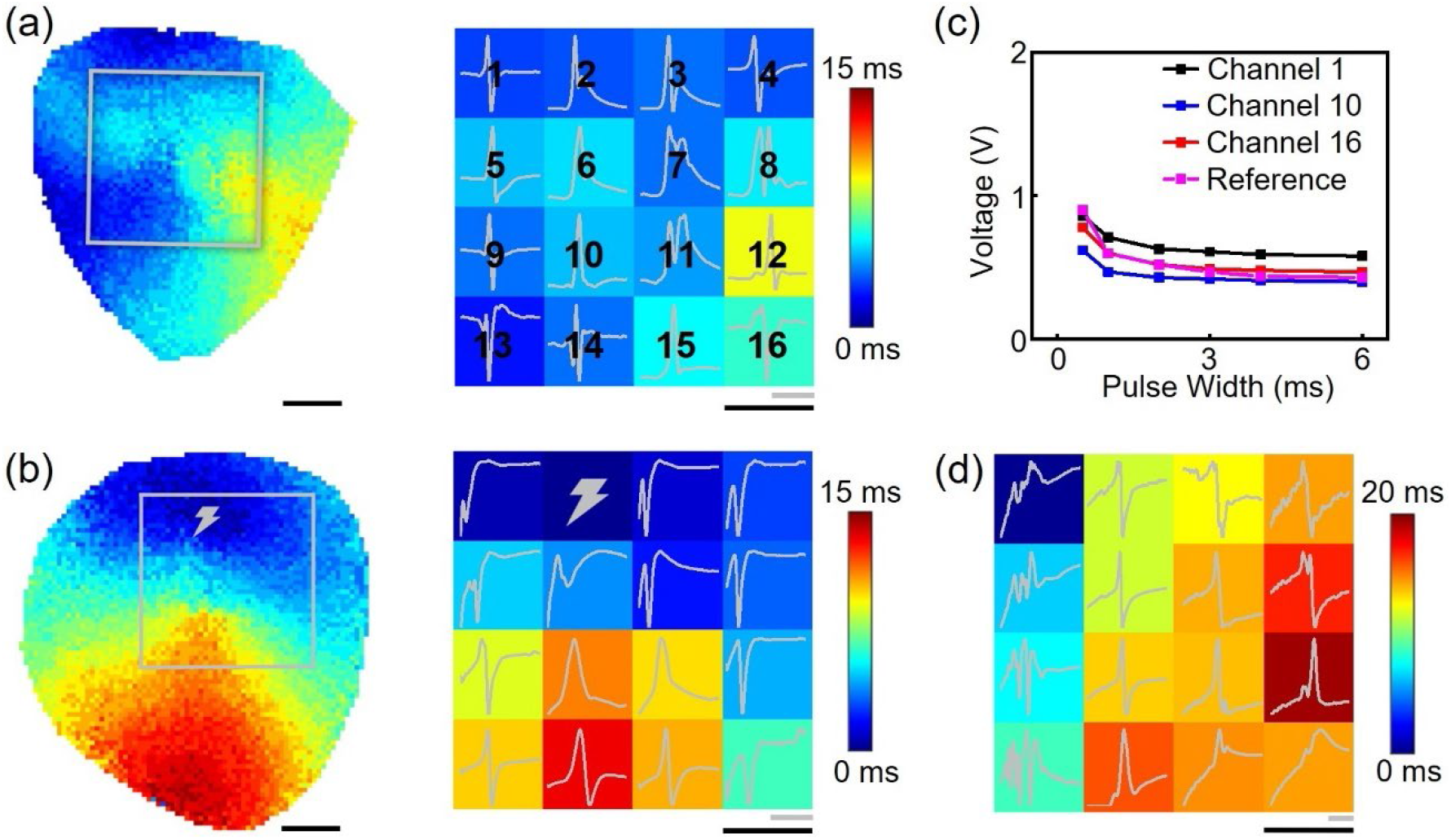
*Ex vivo* demonstration of the electro-optical array devices for spatiotemporal electrical mapping and pacing of mouse hearts and human ventricular tissue slices. Optical (left) and electrical (right) activation maps during sinus rhythm (a) and under pacing (b) from microelectrode channel 2 (indicated by the gray lightning bolt) in the MEA. The 16 microelectrodes are labeled in the electrical activation map in (a). c) Electrical strength duration curves obtained from a heart with microelectrode channels 1, 10, and 16 performing unipolar pacing. Reference strength duration curve is obtained with the Pt-Ir electrode placed at microelectrode channel 16 site. d) Electrical activation map from a human ventricular tissue slice. Gray scale bar, 15 ms. Black scale bar, 1 mm.

### 2.4. *Ex Vivo* Cardiac Electrical Mapping and Optogenetics Pacing

Langendorff-perfused mouse hearts with ChR2-expressing cardiomyocytes in the myocardium serve as a proof-of-concept model to evaluate the capability of the electro-optical array devices in performing bidirectional spatiotemporal electrical mapping and optogenetics pacing. Unlike conventional opaque microelectrodes that block the field of view at the microelectrode sites,^[41]^the Au grid transparent MEA allows colocalized optogenetics interrogation of cardiac activity with the underneath μ-LEDs due to its high optical transparency. **Figure 5**a summarizes the optical strength duration curves (n=3), reflecting the optical irradiance thresholds for successful optogenetics ventricular pacing with a rheobase and chronaxie of 3.91 ± 0.20 mW/mm^2^ and 1.32 ± 0.33 ms, respectively. VT is a life-threatening fast heart rhythm that frequently occurs in patients with myocardial infarction and myocarditis, which can lead to sudden cardiac death. The electro-optical array devices allow concurrent electrical mapping and optogenetics termination of VT (Figure 5b). Here, VT is introduced by our previously reported protocols,^[42–43]^ electrically mapped and confirmed by the MEA. Acute ventricular optogenetics pacing from the μ-LED underneath microelectrode channel 1 at 7 Hz, 1% duty cycle, and 25.3 mW/mm^2^ for 2.86 s successfully terminates VT, which is confirmed by the continuous mapping of the spatiotemporal EG signals during and after optogenetics pacing with the MEA. The heart returns to sinus rhythm after pacing, suggesting no structure or functional damage induced by the optogenetics pacing protocol. The electrical and optical activation maps during VT and optogenetics pacing show consistent morphology (Figure 5b). The activation maps clearly show the anisotropic propagation of the cardiac waves originated precisely from the optogenetics pacing μ-LED site. Those results confirm the localized light delivery to the small tissue volume above the μ-LED. The extracted conduction velocities from the MEA mapping results are 24.3 cm/s and 29.8 cm/s in the transverse and longitudinal directions during VT while these values increase to 27.3 cm/s and 56.3 cm/s during optogenetics pacing. Representative time-aligned EG signals and optical action potentials from the optogenetics pacing site and the farthest diagonal site (Figure S14, Supporting Information) highlight the capability of the MEA in artifact-free detecting fast cardiac signals and differentiating small cardiac conduction differences at different locations under simultaneous optogenetics pacing.

**Figure 5.**
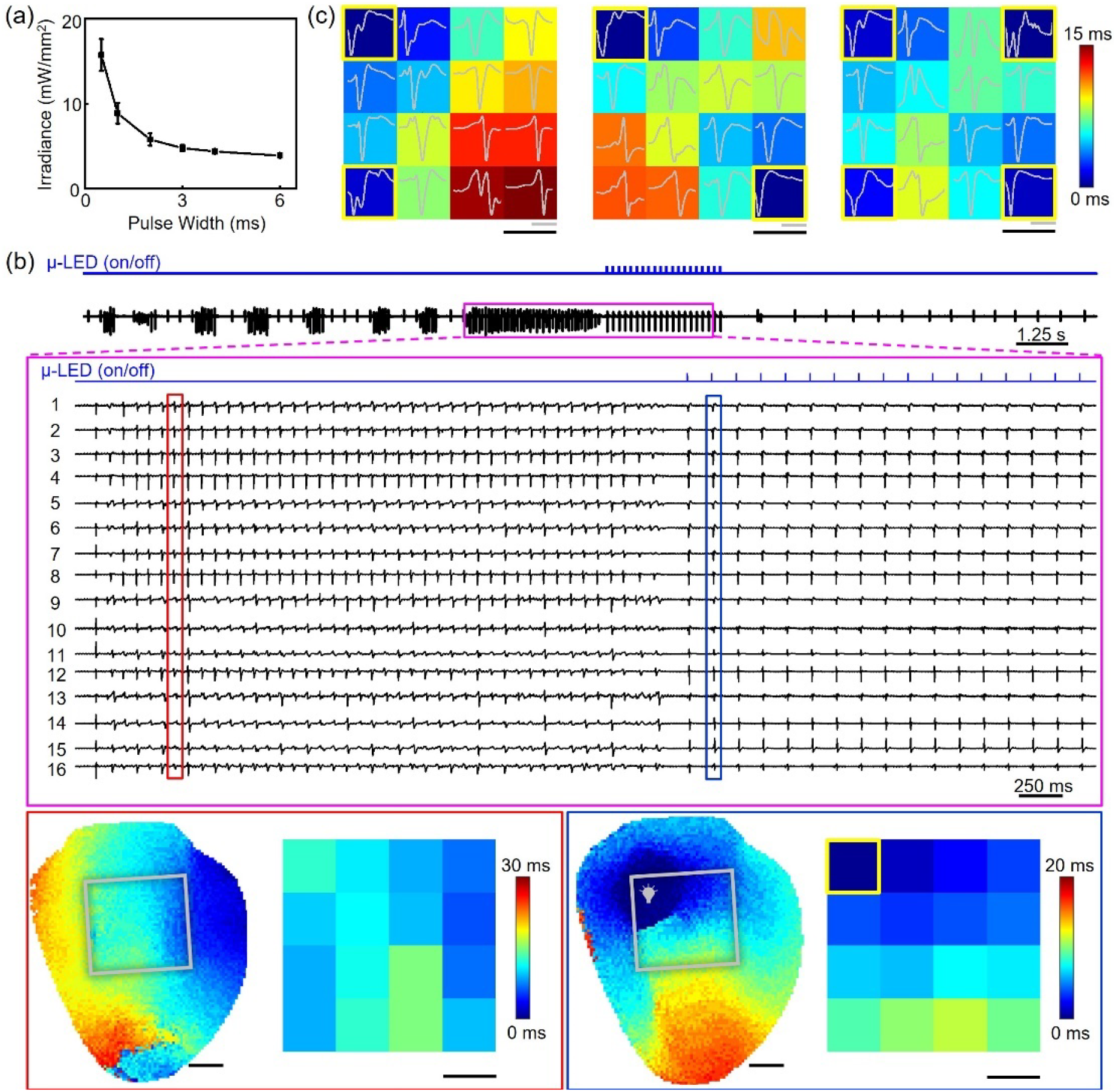
*Ex vivo* demonstration of the electro-optical array devices for spatiotemporal electrical mapping and optogenetics modulation of ChR2-expressing mouse hearts. a) Optical strength duration curves (n=3) obtained from ChR2-expressing mouse hearts. b) 16-channel EG recording of VT followed by optogenetics pacing (7 Hz, 1% duty cycle, 25.3 mW/mm^2^) to restore sinus rhythm. Red and blue insets show optical and electrical activation maps of cardiac activity during VT and optogenetics pacing. c) Electrical activation maps under optogenetics pacing (5 Hz, 1% duty cycle, 8.32 mW/mm^2^) with multiple μ-LEDs in the electro-optical array device. Sites of pacing are indicated by the gray light bulb or yellow box in the activation maps. Gray scale bar, 10 ms. Black scale bar, 1 mm.

In addition to single site pacing, multisite pacing is critically important to fine control and optimize synchrony of excitation, a strategy that is clinically used for resynchronization therapy to treat patients with conduction delay and symptomatic heart failure.^[21]^ Figure 5c shows the results of optogenetics pacing with 2 neighboring, 2 diagonal, and all 4 μ-LEDs operating at 5 Hz, 1% duty cycle, 8.32 mW/mm^2^, and the simultaneous electrical mapping of the cardiac wave propagation with the electro-optical array device placed on the LV. Our device design enables the precise control and monitoring of the wave direction, velocity, activation time range by adjusting the numbers and positions of the operating μ-LEDs. For example, the measured activation time range reduces from 14.7 ms (Figure 5c, left) to 11.8 ms (Figure 5c, middle), and finally to 8.70 ms (Figure 5c, right) due to the fusion of activation wavefronts generated at the sites of optogenetics pacing.

### 2.5. *In Vivo* Epicardial Electrical Mapping and Optogenetics Pacing

Compared to *ex vivo* experiments, *in vivo* studies better demonstrate the dynamic physiology of the blood-perfused innervated intact beating hearts. The electro-optical array device design allows simultaneous spatiotemporal mapping and modulation of cardiac activity from multiple heart regions with high precision. **Figure 6**a presents an open-chest view of a device attached to both the left atrium (LA) and LV of a mouse heart as a temporary cardiac implant. Representative time-aligned EG traces from microelectrode channel 1 and far-field ECG recording results during sinus rhythm are shown in Figure 6b. The average heart rate (339 ± 1.51 BPM) and PR interval (68.8 ± 2.99 ms) from the microelectrode match well with those from the reference ECG recording (339 ± 1.56 BPM and 70.6 ± 3.04 ms, respectively). Figure 6c,d displays electrical activation maps from the MEA during sinus rhythm and optogenetics pacing with the blue μ-LED underneath microelectrode channel 13. During sinus rhythm, ventricular activation initiates from the sinoatrial node and synchronously propagates through the ventricular specialized conduction system (Purkinje network) whereas during optogenetics pacing, activation is controlled to initiate from the ventricular cardiomyocytes at the pacing sites and does not travel in the fastest pathway for conduction.^[44]^ As a result, the MEA measured conduction velocities in the transverse and longitudinal directions decrease from 105 cm/s and 189 cm/s at sinus rhythm to 27.6 cm/s and 36.3 cm/s during optogenetics pacing, respectively. Figure 6e shows 16-channel high-fidelity electrical mapping results during sinus rhythm and 10 Hz optogenetics pacing. The optically paced EG signals exhibit a high SNR of 34.4 ± 8.71 dB. Figure 6f,g demonstrates the spatiotemporal voltage maps at 5 sequential time points of sinus rhythm and optogenetics pacing, highlighting the control and capture of the different activation and propagation profiles of the cardiac excitation waves with the electro-optical array device. During optogenetics pacing, the activation originates from the bottom left area where the pacing μ-LED is located and the anisotropic signal propagation starts from the left of the device to the right regions. Figure S15 (Supporting Information) shows the detection of heart rates during sinus rhythm and optogenetics pacing at a different frequency (8 Hz). The devices further allow *in vivo* programmable single site pacing from different locations (Figure 6h) or multisite pacing (Figure 6i) to realize dynamic spatiotemporal control of the cardiac excitation waves and their propagations. Similar to the *ex vivo* results in Figure 5c, *in vivo* multisite optogenetics pacing results in a reduced activation time range (9.45 ms) than single site pacing (13.4 ms).

**Figure 6.**
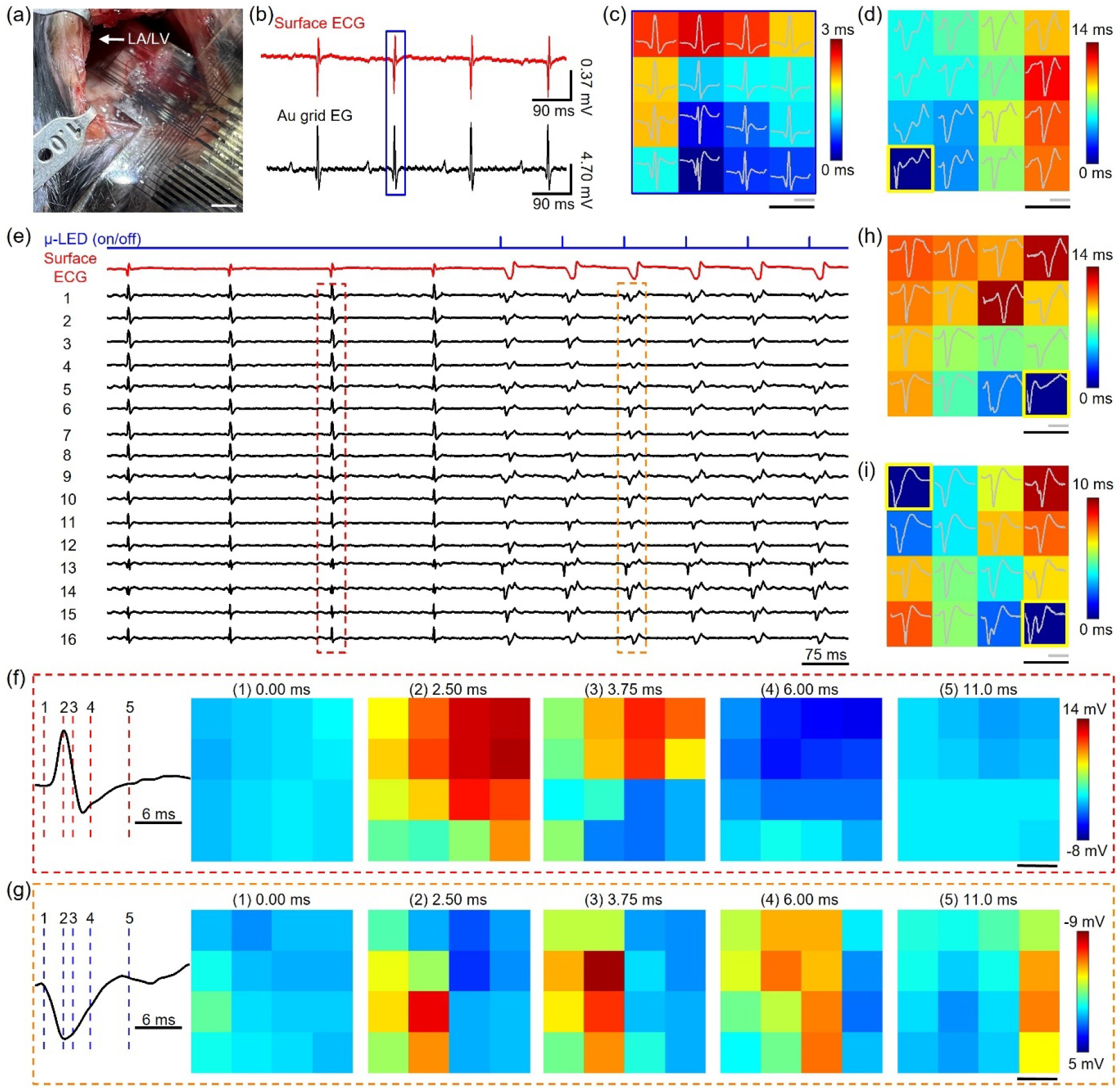
*In vivo* demonstration of the electro-optical array devices for simultaneous spatiotemporal electrical mapping and optogenetics modulation. a) Optical image of the electro-optical array device contacting the LA and LV of a mouse heart *in vivo*. Scale bar, 2 mm. b) Surface ECG and EG from microelectrode channel 1 in the Au grid transparent MEA show high-fidelity recording of atrial and ventricular activity during sinus rhythm. c) Electrical activation map recorded by the MEA during sinus rhythm at the time window highlighted by the blue box in (b). d) Electrical activation map under single site optogenetics pacing (10 Hz, 5% duty cycle, 14.0 mW/mm^2^). e) 16-channel EG mapping results during sinus rhythm and optogenetics pacing using the pacing condition in (d). Voltage maps from EGs in (e) showing 5 sequential time points during sinus rhythm (f, red dashed box) and optogenetics pacing (g, orange dashed box). Electrical activation maps during single site (h) and multisite (i) optogenetics pacing. Sites of optogenetics pacing are indicated by the yellow box in the activation maps. Gray scale bar, 15 ms. Black scale bar, 1 mm.

## 3. Conclusion

In conclusion, we have achieved soft electro-optical array devices integrating multilayered transparent MEAs directly on top of multicolor μ-LED arrays in thin, flexible platforms for mechanically compliant, multiplexed, spatiotemporal cardiac electrical mapping/pacing and optogenetics modulation *ex vivo* and *in vivo*. Those devices directly address the limitations from conventional optical fiber based implantable cardiac optogenetics tools that lack spatiotemporal sensing capabilities and exhibit limited optical channel densities for multisite and/or multicolor actuation and control of heart rhythm. The transparent MEA provides high-fidelity regional mapping of cardiac EG signals under sinus rhythm, clinically relevant complicated cardiac conditions, such as VT, and optogenetics pacing. The devices show single-/multi-site pacing capabilities with differing parameters for dynamic control of the spatiotemporal cardiac activity, both electrically and optically, to study the efficacy of various cardiac pacing strategies on treating arrhythmias and heart failure. The device features are crucial in (1) artifact-free measuring the real-time cardiac rhythm, such as excitation patterns and electrical wave propagations, before, during, and immediately after the colocalized site-specific optogenetics pacing; (2) investigating the spatiotemporal responses of and the underlying mechanisms involved in different optogenetics pacing therapies; (3) unraveling the different effects from electrical and optogenetics pacing. The multifunctional operation modes have great potential for automated closed-loop operations, whereby cardiac activity can be altered using patterned illumination based on the incoming activity being measured with the MEA.

Device fabrication and integration (e.g., micro-transfer printing, photolithography, etc.) are built upon scalable processes where the number of channels and the pitch values can be scaled up or customized based on the specific needs (e.g., for applications in large animal models, improve the mapping resolution). The tunable emission profiles of the μ-LEDs allow flexibility for the inclusion of additional optogenetics actuators and validating their effectiveness for cardiac research. Our current devices are designed as a temporary epicardial implant. Future work should focus on designing wireless power supply, control/communication,^[45–46]^ and strategies to minimize foreign-body responses^[47–48]^ for realizing chronic, fully implantable, multifunctional cardiac implants. Those approaches would allow us to better identify and understand the long-term abnormalities in cardiac electrophysiological function that occur over the course of *in vivo* heart disease development, progression, treatment, and potentially advance the discovery of therapeutic treatment of lethal heart disease.

## 4. Experimental Section

### Device Fabrication

The fabrication of the device began with the lamination of a 25 μm thick PET film on a PDMS-coated glass slide followed by spin-coating of a 7 μm thick SU-8 (Kayaku Advanced Materials Inc.) planarization layer. Next, μ-LED interconnects and bonding pads were lithographically defined using AZ® nLOF 2070 (Integrated Micro Materials) photoresist, followed by electron beam (E-beam) evaporation of 20/100 nm of Cr/Cu and liftoff in acetone. Blue and green μ-LEDs (C460TR2227/C527TR2227, Cree Inc.) were micro-transfer printed using a soft lithography defined PDMS array stamp onto μ-LED bonding pads, followed by a reflow soldering process at 150 °C with an In/Ag solder paste (Indalloy 4, Indium Corporation) to form robust electrical and mechanical connections. The μ-LEDs were insulated by a 60 μm thick SU-8 layer. The Au grid transparent MEA and interconnect patterns were lithographically defined on the insulation SU-8 above the μ-LEDs, followed by E-beam evaporation of 20/40 nm of Cr/Au and liftoff in acetone. Another 7 μm thick SU-8 layer encapsulated the Au grid transparent MEA and defined the microelectrode windows. The device was connected out to a printed circuit board that interfaced with external power supply and data acquisition system using a flexible anisotropic conductive film (ACF) (Elform). The interface between the bonding pads on the device and the ACF cable was coated with PDMS to prevent current leakage into the biofluids during measurement.

### Optical, mechanical, electrical, and thermal characterizations

A spectrometer (AvaSpec-ULS2048L) measured the emission profiles and irradiances of the multicolor μ-LEDs. The blue light attenuation profile on the mouse epicardial surface was obtained from a MiCAM05 high-speed CMOS imaging system (SciMedia). A custom MATLAB program was used to analyze the optical propagation profile of the emission light. Transmittance of the Au grids was measured using a spectrophotometer (V-770, Jasco Inc.). A motorized test stand (ESM 1500, Mark-10) assessed the mechanical performance of the electro-optical array devices. A Keithley 2614B source meter and probe station measured the current-voltage characteristics of the μ- LEDs. A FLIR E8-XT thermal camera monitored the local temperature changes of the devices with the μ-LEDs operating at different conditions.

### Electrochemical characterizations

EIS and CV studies were conducted using a potentiostat (Gamry Reference 600+ potentiostat/galvanostat/ZRA, Gamry Instruments Inc.) in a three-electrode configuration containing a Ag/AgCl reference electrode, a Pt counter electrode, and a Au grid microelectrode serving as the working electrode. All microelectrode samples were plasma cleaned for 1 minute at 100 W and then measured in a 1× PBS solution (Sigma-Aldrich). Frequencies were swept from 100 Hz to 10 kHz with a 10 mV RMS AC voltage for EIS studies. A bioamplifier (ADInstruments) provided 0.5 V cathodal electrical pulses at 1.6% duty cycle and 8 Hz frequency to stimulate the microelectrodes. CSC was calculated using a 100 mV/s scan rate. A Pt electrode connected to a PowerLab data acquisition system (ADInstruments) input a 20 mV peak-to-peak sine wave into the 1× PBS solution for the benchtop electrical recording characterization of the Au grid microelectrodes. The microelectrodes detected the sinusoidal signals and output the signals to the PowerLab data acquisition system where they were filtered by 50 Hz notch filters and a 0.3 Hz to 2 kHz bandpass filter. The signals were subsequently processed in a custom MATLAB program for analysis of SNR and RMS noise.

### Mouse model

All animal procedures were in accordance with the ethical guidelines of the National Institutes of Health and approved by the Institutional Animal Care and Use Committee of the George Washington University. Wild-type C57BL/6 mice (Jackson Labs) were used for experiments assessing the electrical recording and stimulation capabilities of the MEAs. For experiments involving optogenetics, we used commercially available transgenic mice and crossbred them as previously described^[45]^ to express ChR2 in myocardial cells.

### Ex vivo device demonstration

For *ex vivo* demonstrations, mice were anesthetized through the inhalation of a mixture of 3% isoflurane vapor and 97% O_2_. The cessation of pain was verified by a toe pinch followed by an open thoracotomy to quickly excise the heart. Upon excision, the heart was cannulated and Langendorff-perfused with oxygenated (95% O_2_ and 5% CO_2_) Tyrode’s solution (37 °C) under an aortic pressure from 60 mmHg to 80 mmHg. 15 μM excitation-contraction uncoupler blebbistatin (Cayman Chemical) was used to arrest heart contraction and avoid motion artifacts. The electro-optical array device was then placed against the epicardial surface of the LV to record EG signals that output to a 16-channel PowerLab data acquisition system at 20 kHz sampling frequency. The signals were filtered with a 0.5 Hz to 2 kHz bandpass filter and analyzed in a custom MATLAB program. A current source (Keithley 6221) operated the μ-LEDs in the electro-optical array device. Optical fluorescence mapping was conducted with a high-speed MiCAM05 CMOS camera using potentiometric dyes RH237 or di-4-ANBDQBS for wild-type and ChR2-expressing mouse hearts, respectively. The fluorescence signals were filtered with long-pass filters. The optical mapping results were analyzed using the custom MATLAB software (RHYTHM) that is available at https://github.com/optocardiography. VT was induced via ischemic reperfusion until confirmation by the electro-optical array device, as described previously.^[42–43]^ Electrical and optical strength duration curves were obtained at pulse widths of 0.5 ms, 1 ms, 2 ms, 3 ms, 4 ms, and 6 ms, respectively. 1:1 capture was observed for 10 seconds before confirming capture at a specified pulse width.

### In vivo device demonstration

Mice under surgery were intubated using a 20 gauge endotracheal tube and ventilated using a VentElite small animal ventilator (Harvard Apparatus). Mice were anesthetized using a mixture of 3% isoflurane vapor and 97% O_2_. They were administered a dose of buprenorphine (subcutaneously) at a dosage of 0.05-0.10 mg/kg in 0.25 mL of saline. Pain cessation was verified by a toe pinch after decreasing isoflurane to 2% before the onset of surgical incisions to open the thoracic cavity. The electro-optical array device was then placed inside the thoracic cavity onto the epicardial surface of the LA and LV. Unipolar EG signals were recorded at a 20 kHz sampling frequency and filtered with 0.5 Hz to 2 kHz bandpass filter. A custom MATLAB program was used to analyze the obtained signals where the time of activation was calculated using the minimum derivative of the time-domain signal. A three-electrode subcutaneous ECG using a PowerLab data acquisition system served as a reference for all *in vivo* studies.

### Donor Human Heart Experiments

Experiments using donor human heart tissue were approved by the Institutional Review Board at the George Washington University. Donor human hearts were procured from Washington Regional Transplant Community in Washington, DC as deidentified discarded tissue. Heart slices (thickness, 400 μm) were prepared using our previously reported protocol.^[49]^ The tissue was perfused with Tyrode’s solution and treated with blebbistatin to minimize motion artifacts. Voltage-sensitive fluorescent dye di-4-ANBDQBS stained the tissue. The electrical and optical mapping were acquired and analyzed similarly to those in the *ex vivo* and *in vivo* animal experiments above.

## Acknowledgements

We thank the Nanofabrication and Imaging Center at the George Washington University for its facilities regarding device fabrication and characterization. L. L. and I. R. E. acknowledge National Science Foundation grants (2011093 and 2131682). L. L. acknowledges the George Washington University Cross-Disciplinary Research Fund and University Facilitating Fund.

## Author Contributions

S.N.O., M.M., I.R.E., and L.L. conceived the overall research goals and objectives. S.N.O., Z.C., J.T., C.H., J.A., N.D., and J.B. performed device fabrication, characterization, and analysis. S.N.O., M.M., and Z.L. performed *ex vivo* and *in vivo* experiments and data analysis. S.N.O., M.M., I.R.E., and L.L. wrote the manuscript.

