## Supplementary Material for "Multifunctional flexible electro-optical arrays for simultaneous spatiotemporal cardiac mapping and modulation"

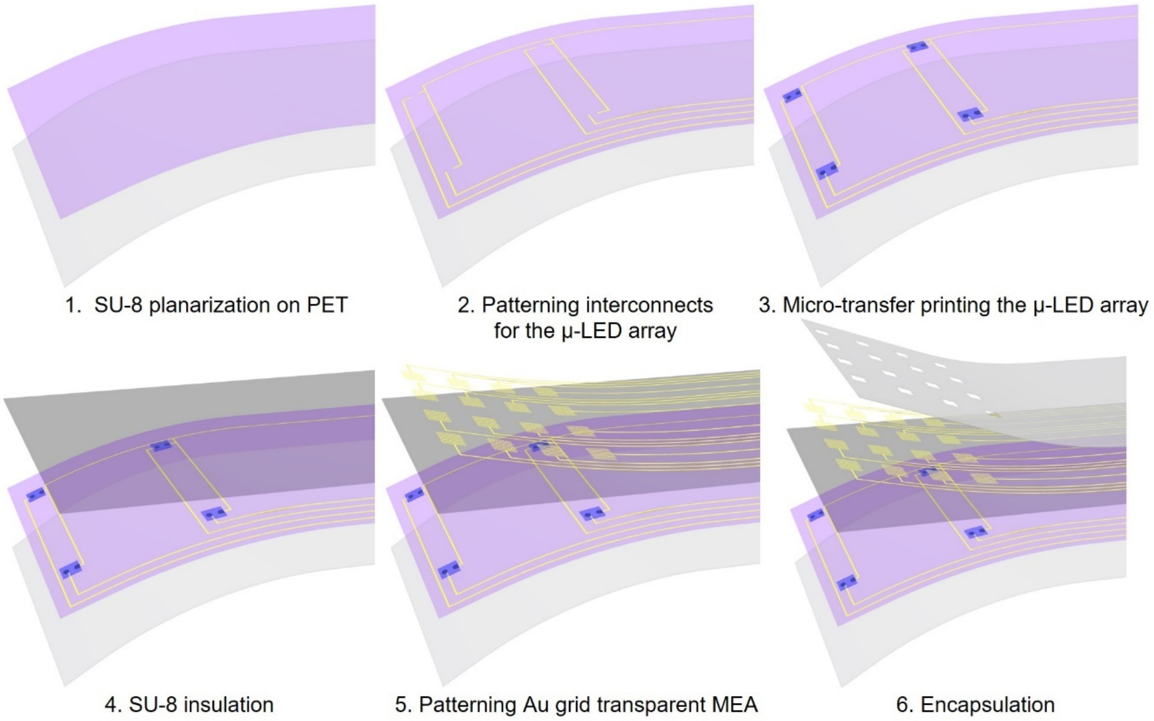


**Figure S1.** Fabrication scheme of the multifunctional electro-optical array device.

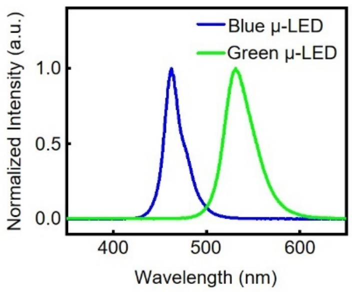


**Figure S2.** Emission spectra of the blue and green µ-LEDs used in the multifunctional electro-optical array devices.

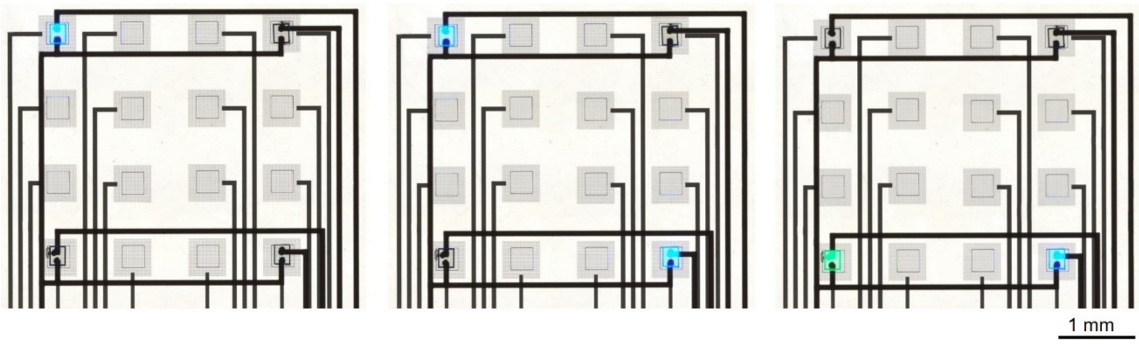


**Figure S3.** Optical images of the electro-optical array devices. From left to right: devices with 1 blue μ-LED on, 2 blue μ-LEDs on, 1 blue and 1 green μ-LEDs on, respectively.

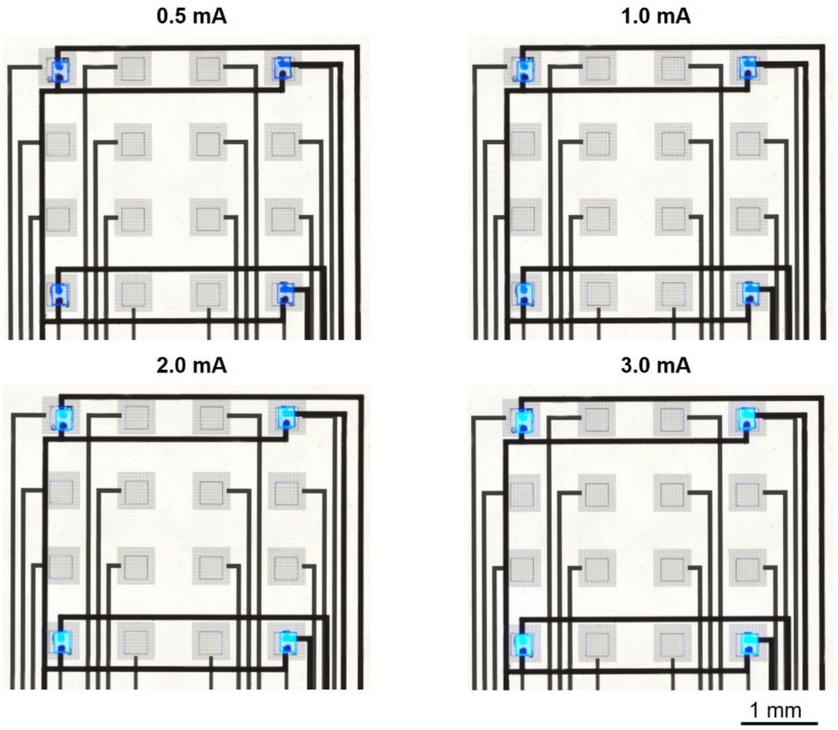


**Figure S4.** Optical images of the electro-optical array devices with the 4 blue μ-LEDs operated at varying forward current conditions from 0.5 mA to 3.0 mA.

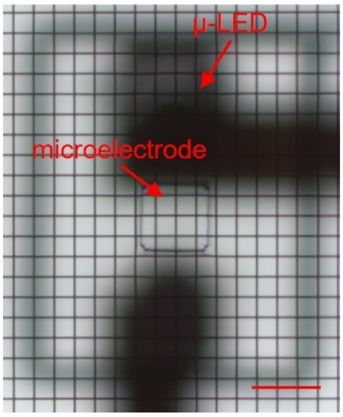


**Figure S5.** An electro-optical array device containing cellular-scale Au grid transparent microelectrodes with dimensions at 50 × 50 $\mu m^2$ in the MEA. Scale bar, 50 μm.

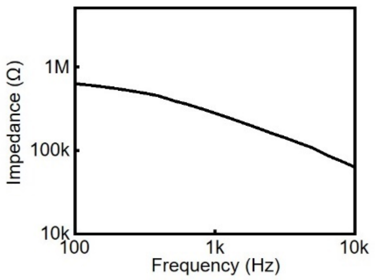


**Figure S6.** Impedance spectra of the 50 × 50 μm$^2$ cellular-scale Au grid transparent microelectrode in the multifunctional array device in Figure S5.

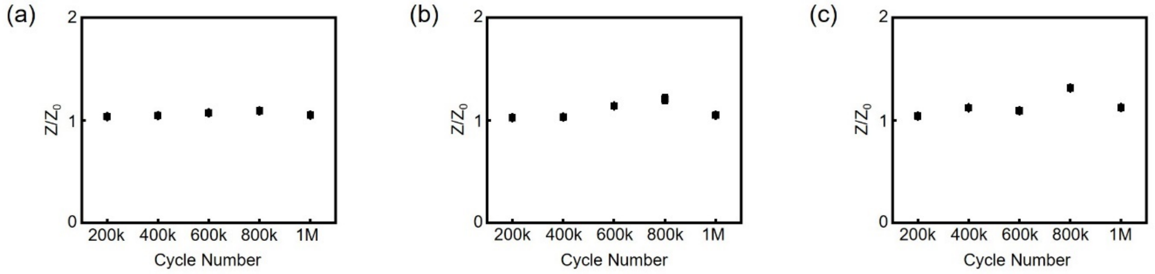


**Figure S7.** (a to c): changes of Au grid transparent microelectrode impedance as a function of electrical stimulation cycle (8 Hz, 1.6% duty cycle, 0.5 V) from three different multifunctional array devices. Z is the impedance at a specific electrical stimulation cycle and $Z_0$ is the initial impedance, respectively.

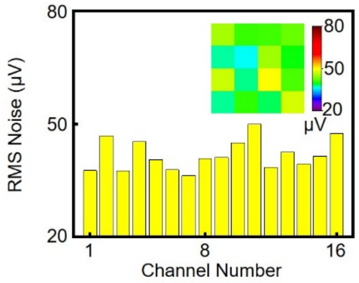


**Figure S8.** RMS noise bar chart of the 10 Hz, 20 mV peak-to-peak input sine wave recorded by the Au grid transparent MEA. Inset: SNR colormap of the 16 microelectrodes with respect to actual microelectrode position in the Au grid transparent MEA.

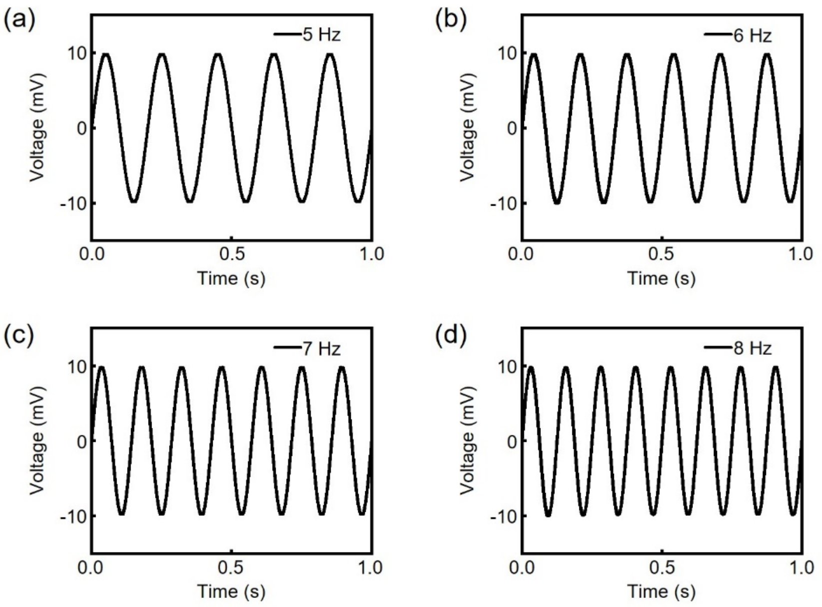


**Figure S9.** Superimposed benchtop electrical recording output of 20 mV peak-to-peak input sine waves from 4 different Au grid transparent microelectrodes at 5 Hz (a), 6 Hz (b), 7 Hz (c), and 8 Hz (d).

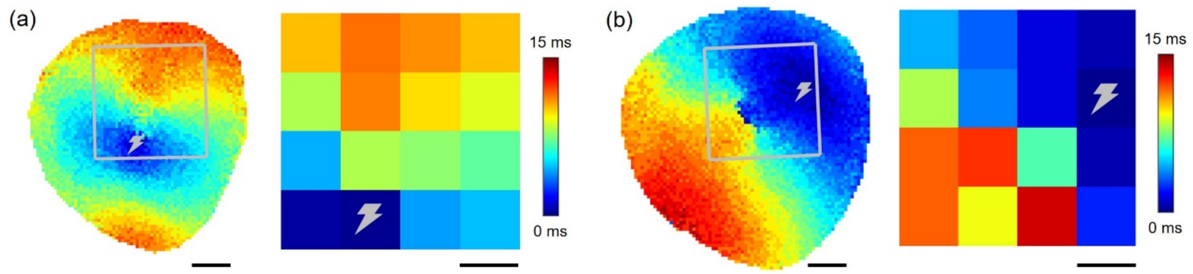


**Figure S10.** Optical activation maps (left) and device electrical activation maps (right) obtained from a heart under electrical stimulation from microelectrode channel 14 (a) and channel 8 (b). Sites of electrical stimulation are indicated by the gray lightning bolt.  Scale bar, 1 mm.

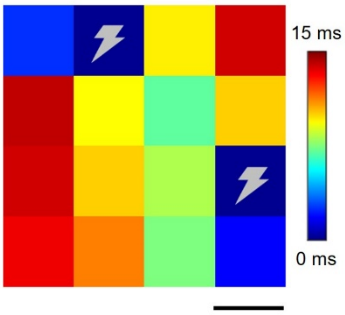


**Figure S11.** Electrical activation map obtained with the Au grid transparent MEA from a mouse heart under two sites electrical stimulation from microelectrode channels 2 and 12. Sites of stimulation are indicated by the gray lightning bolt. Scale bar, 1 mm.

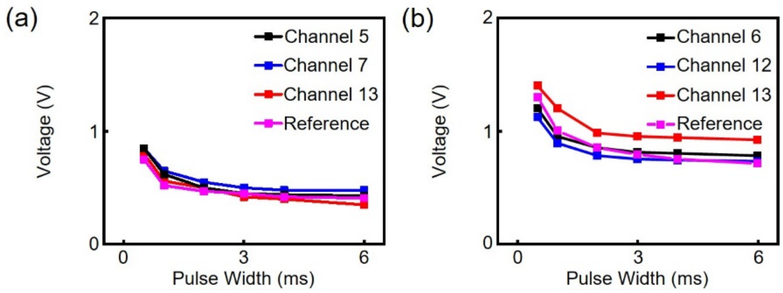


**Figure S12.** Strength duration curves obtained with different microelectrode channels in a Au grid transparent MEA from mouse 1 (a) and mouse 2 (b).

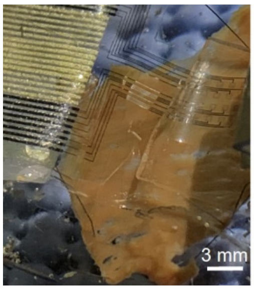


**Figure S13.** Optical image of a Au grid transparent MEA positioned on a human ventricular tissue slice for spatiotemporal electrical and optical mapping of cardiac wave propagations.

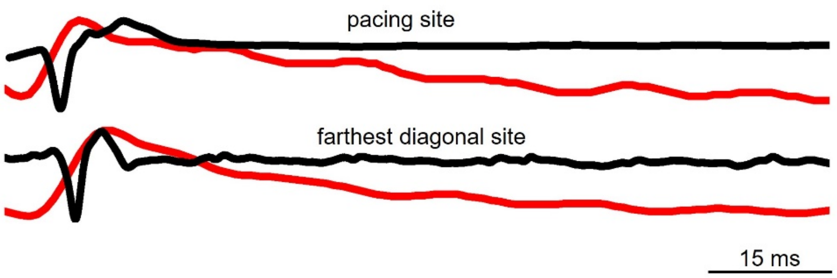


**Figure S14.** Time-aligned EG signals (black) and optical mapping signals (red) from the microelectrode channels at the optogenetics pacing site and the farthest diagonal site.

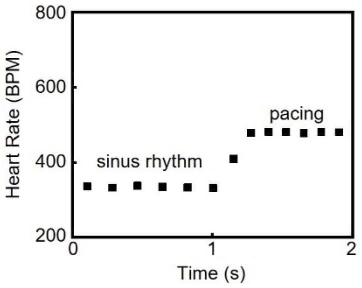


**Figure S15.** *In vivo* heart rates recorded by the Au grid transparent MEA before and during single site optogenetics pacing at 8 Hz, 14.0 mW/mm$^2$, and 5% duty cycle.
